# Effects of Phthalate Metabolite Mixture Exposure on Mouse Oocyte Development

**DOI:** 10.64898/2026.05.20.726577

**Authors:** Juan Dong, Vidhi Patel, Shuangqi Wang, Hasanur Alam, Wenjie Yang, Anika Roy, Leyi Wang, Jodi A. Flaws, Huanyu Qiao

## Abstract

Phthalates are pervasive endocrine-disrupting chemicals widely used in consumer products. The wide use of many phthalates results in chronic human exposure to complex mixtures rather than single compounds. Despite extensive studies on individual compounds, the combined effects of phthalate metabolites on oogenesis remain poorly understood. Here, we developed a precise microinjection-based single-oocyte toxicological assay to examine the impact of a defined phthalate metabolite mixture on meiotic progression. Phthalate mixture exposure markedly impaired oocyte maturation, as most oocytes failed to extrude the first polar body. Mechanistic analyses revealed severe meiotic defects, including disrupted spindle morphology, chromosome misalignment, disorganized actin cytoskeleton, and impaired mitochondrial function, accompanied by excessive reactive oxygen species (ROS) accumulation and DNA damage. Single-cell transcriptomic profiling further identified differentially expressed genes enriched in biological processes related to exocytosis, secretory pathway regulation, and cytoskeletal organization, as well as in MAPK, JAK-STAT, cGMP-PKG, and GnRH signaling pathways that are essential for follicular development and oocyte maturation. Together, these findings demonstrate that combined phthalate exposure directly compromises female gamete quality and underscore the importance of evaluating mixture effects when assessing risks to women’s reproductive health.

## Introduction

Infertility, defined as the failure to achieve a clinical pregnancy after 12 months of regular, unprotected intercourse, is a major global health issue, with the lifetime prevalence of infertility estimated to be approximately 17.5% worldwide [1]. In recent years, the incidence of female infertility risen steadily, largely due to a decline in both the quantity and quality of oocytes [2]. In adult female mammals, fully grown oocytes are arrested at the diplotene stage of meiotic prophase I, known as the germinal vesicle (GV) stage [3]. In humans, this arrest may persist for decades [4]. Upon luteinizing hormone (LH) stimulation, meiosis resumes with germinal vesicle breakdown (GVBD). Following GVBD, microtubules reorganize to assemble the meiotic spindle near the oocyte nucleus, which subsequently migrates toward the cortex by the end of metaphase I (MI). This spindle translocation, driven primarily by the actin cytoskeleton, is essential for establishing asymmetrical division required for polar body extrusion. Once the spindle reaches the cortex, chromosomes align properly on the metaphase plate [5, 6]. Meiotic progression proceeds to anaphase I (AI), during which homologous chromosomes segregate and the cytoplasm divides asymmetrically, resulting in the extrusion of a small polar body and formation of a mature oocyte [7]. Importantly, mammalian oocyte meiosis is highly sensitive to environmental insults, and exposure to toxic agents can cause irreversible damage with long-term consequences for developmental competence [8].

Phthalates, a major class of endocrine-disrupting chemicals (EDCs), are commonly used as plasticizers and solvents in various consumer products, including food packaging, building materials, personal care products, and medical devices [9, 10]. Since phthalates are not chemically bound to the materials in which they are incorporated, they can easily leach into the environment and can enter the human body through ingestion, inhalation, or dermal absorption [9, 11]. Once absorbed, phthalate diesters are rapidly hydrolyzed to monoester metabolites, which may undergo further oxidization to secondary products and distribute broadly across tissues [12]. Thus, phthalate metabolites have been widely detected in various human body fluids, including serum, breast milk, sweat, urine, and follicular fluid [13–17], and monoester forms are generally considered more toxic than their parent compounds [18–21].

Growing evidence links phthalate exposure to adverse reproductive outcomes in animal models and women [22]. These adverse outcomes include infertility, pregnancy loss, congenital abnormalities, and impaired offspring health phthalate-induced oocyte defects linked to infertility being the primary focus of the current study. Phthalates metabolites have been detected in human follicular fluid, indicating that these compounds can directly reach and affect oocytes [15]. As EDCs, phthalates disrupt ovarian function by inhibiting follicle growth, impairing oocyte development, reducing steroid hormone production, and promoting follicular atresia [20, 23]. Exposure to the individual phthalate, di(2-ethylhexyl) phthalate (DEHP), at environmentally relevant levels (approximately 100 µmol/L) increases meiotic double-strand breaks (DSBs) and impairs chromosome remodeling in oocytes [24]. DEHP also disrupts antral follicle growth by dysregulating pathways involved in the cell cycle, apoptosis, and steroidogenesis, and oxidative stress [20, 25], and it impairs primordial follicle assembly in the neonatal ovary by upregulating phosphodiesterase enzyme 3A (PDE3A) [26]. Similarly, dibutyl phthalate (DBP) induces oxidative stress, DNA damage, and downregulation of key meiotic regulators, ultimately triggering oocyte apoptosis [27]. Collectively, these findings demonstrate that single phthalate exposure can profoundly disrupt ovarian physiology, thereby posing a significant risk to female fertility.

Humans are exposed to complex mixtures of phthalates rather than isolated compounds daily, but research on phthalate mixtures remains limited. In previous studies, an environmentally relevant phthalate mixture (21% DEHP, 35% diethyl phthalate (DEP), 15% DBP, 8% diisobutyl phthalate (DIBP), 5% benzylbutyl phthalate (BBzP), and 15% di-isononyl phthalate (DINP)) disrupted follicle development, compromised hormone synthesis, and increased follicular atresia in cultured antral ovarian follicles [28]. The same mixture has been shown to suppress ovulation by modifying progesterone pathway activity and extracellular matrix remodeling [29]. More recently, neonatal exposure to this environmentally relevant phthalate mixture was shown to reduce primordial follicle number and disrupt steroidogenic, apoptotic, and inflammatory pathways [30]. In contrast, a monoester phthalate mixture (37% monoethyl phthalate (MEP), 19% mono(2-ethylhexyl) phthalate (MEHP), 15% monobutyl phthalate (MBP), 10% monoisononyl phthalate (MNP), 10% monoisobutyl phthalate (MiBP), and 8% monobenzyl phthalate (MBzP)), reflecting relative proportions of metabolites detected in the urine of women and the metabolites that reach the ovary and are present in follicular fluid, increases expression of cell-cycle inhibitors and anti-apoptotic factors in neonatal ovarian cultures [31]. Epidemiological studies further show that multiple phthalate metabolites are found in human follicular fluid and that higher phthalate exposure correlates with poorer IVF outcomes [32–34]. These findings highlight the urgent need to elucidate the mechanistic effects of phthalate mixtures on oocyte quality.

In the present study, we elucidated the adverse effects of a phthalate metabolite mixture on mouse oocytes using a direct microinjection approach. Key indicators of oocyte quality, including first polar body extrusion, cytoskeletal organization, and mitochondrial function, were systematically examined. The concentration of the phthalate metabolite mixture was selected to reflect levels detected in human follicular fluid [33–35], thereby ensuring physiologically relevant exposure conditions. To our knowledge, this is the first study to directly assess the effects of a phthalate mixture on oocytes. Furthermore, single-cell transcriptomic analysis was conducted to delineate the molecular pathways underlying phthalate mixture-induced disruptions in oocyte maturation and developmental competence.

## Methods

### Chemicals

The phthalate mixture was formulated to reflect human-relevant exposure and contained 36.7% MEP, 19.4% MEHP, 15.3% MBP, 10.2% MiBP, 10.2% MiNP, and 8.2% MBzP by weight. The relative proportions were determined based on the molar sums of these metabolites measured in the urine of pregnant women [36]. Importantly, these phthalate metabolites are widely detected in women across different age groups and geographic regions, including population-based studies conducted in the United States and other countries [37–40]. Similar metabolites have also been identified in human follicular fluid and ovarian samples [41], supporting the biological and environmental relevance of the selected metabolites A final concentration of 40 nM was selected for microinjection in this study, which is markedly lower than that used in previous follicle or ovarian culture studies involving individual phthalates or phthalate mixtures [31, 42, 43]. This concentration was chosen to reflect the physiological levels observed in women’s follicular fluid. Numerous studies have reported the presence of phthalate metabolite mixtures in human follicular fluid, with concentrations generally in the ng/mL range [33–35]. It is well established that urine is the preferred matrix for assessing total daily phthalate exposure, as these non-persistent compounds are rapidly metabolized and excreted, resulting in higher and more representative urinary levels than in serum [12]. However, the weak to moderate correlations between urinary and tissue (particularly follicular fluid) concentrations indicate that urinary levels may not accurately reflect exposure in ovarian or other reproductive tissues [44]. Therefore, follicular fluid represents a more appropriate matrix for evaluating ovarian-specific effects. Importantly, since this study employed direct microinjection of the phthalate metabolite mixture into oocytes, it focuses on the effects of phthalates under direct exposure conditions rather than through follicle or ovary culture. Accordingly, using concentrations comparable to those found in follicular fluid provides a physiologically relevant reference for determining the injection dose. The concentration of each component in the mixture falls within this physiological range, ensuring that the experimental conditions closely mimic environmentally relevant human exposure. Detailed information on the mixture composition and concentrations is provided in Supplemental Table 1 and 2.

### Animals

CD-1 mice (Charles River Laboratories, Wilmington, MA) were maintained in the Animal Care Facility of the University of Illinois Urbana-Champaign (UIUC) under controlled environmental conditions (12 h light/12 h dark cycle, 22 ± 1 °C) with free access to food and water. All animal procedures were conducted in accordance with protocols approved by the Institutional Animal Care and Use Committee (IACUC) of UIUC.

### Cumulus-oocyte-complexes (COCs) collection and *in vitro* maturation

Ovaries were collected from 4–6-week-old female mice following euthanasia, washed in PBS, and transferred to pre-warmed (37 °C) M2 medium (Sigma, USA). Cumulus–oocyte complexes (COCs) were released by puncturing antral follicles with sterile needles and collected into fresh M2 droplets. The COCs were microinjected with either water or phthalate metabolite mixtures, washed three times, and transferred into pre-warmed in vitro maturation (IVM) medium consisting of α-minimum essential medium (α-MEM; Gibco, USA) supplemented with 10% fetal bovine serum (FBS; VWR Seradigm, USA), 200 mIU/mL follicle-stimulating hormone (FSH; Millipore, USA), and 50 µg/mL sodium pyruvate (Corning Cellgro, USA). The injected COCs were cultured at 37 °C in a 5% COL incubator, and oocytes were collected at defined time points corresponding to different meiotic stages. For germinal vesicle (GV) stage analysis, COCs were cultured for 2 h and then denuded by gentle pipetting. For metaphase I (MI) stage, COCs were cultured for 6 h, denuded, and further cultured for 2 h in M16 medium (Sigma, USA). For metaphase II (MII) stage, COCs were cultured for 6 h, denuded, and further cultured for 12 h in M16 medium. The collected oocytes were used for subsequent analyses. At the end of the culture period, the proportions of oocytes with normal polar body extrusion, degenerated oocytes, and unhealthy oocytes were assessed. Unhealthy oocytes were defined as those exhibiting granular cytoplasm, cytoplasmic shrinkage, or detachment of the oocyte membrane from the zona pellucida, resulting in an increased perivitelline space.

### Phthalate metabolite mixture treatment and *in vitro* maturation

Phthalate metabolite mixture was dissolved in dimethyl sulfoxide (DMSO) and diluted in M16 maturation medium to final concentrations of 0.4 μM and 1.4 μM, with DMSO ≤ 0.1% in all treatments. Control oocytes were cultured in M16 containing the same DMSO concentration without the phthalate mixture. GV oocytes were cultured in vitro for 18 h, after which MII-stage oocytes were collected for subsequent experiments.

### Single-cell microinjections

Borosilicate glass capillaries (TW100-4, World Precision Instruments Inc, FL) were pulled into either holding or microinjection pipettes using a horizontal micropipette puller (P-97, Sutter Instrument Co., CA, USA). The holding pipette was used to stabilize cumulus–oocyte complexes (COCs) during microinjection, while the injection pipette delivered the phthalate metabolite mixture into the oocyte cytoplasm. The diluted phthalate metabolite mixture was backfilled into the injection pipette using a microloader tip (100 mm, 20 μL; Fisher Scientific, MA, USA). The holding pipette was mounted on the left-side micropipette holder of a hydraulic micromanipulator, and the injection pipette was mounted on the right-side holder. Both were connected to a FemtoJet 4i microinjector (Calibre Scientific, OH, USA), which controlled injection pressure and duration to ensure precise delivery of the phthalate metabolite mixture.

To determine the injection volume, the diameter of the expelled droplet was measured in mineral oil, and its volume was calculated using the formula V = 4/3 πr³. Under the defined parameters (injection pressure 50 hPa, injection time 0.10 s, and compensation pressure 20 hPa), the average droplet volume was 0.523 pL. Given an average oocyte diameter of 80 µm (calculated volume = 268.083 pL), the stock concentration of the injected phthalate metabolite mixture required to achieve a final intracellular concentration of 40 nM (comparable to follicular fluid levels) was determined using the formula X = 268.083 × target concentration / 0.523, resulting in a calculated stock concentration of 20 µM. Control oocytes were injected with an equivalent volume (0.523 pL) of sterile water under identical conditions.

During microinjection, each COC was positioned with the microinjection pipette gently pressing against the holding pipette to facilitate precise penetration through the zona pellucida. The phthalate metabolite mixture was then delivered into the oocyte cytoplasm by controlling injection pressure and duration. All COCs underwent identical handling and were subsequently cultured in IVM medium for further analysis.

### N-acetyl-L-cysteine (NAC) treatment

Following microinjection with either the phthalate metabolite mixture or water (control), GV-stage oocytes were randomly assigned to three experimental groups. Control and phthalate-exposed oocytes were cultured in pre-warmed IVM medium and subsequently transferred to M16 medium, as described above. For the rescue group, a subset of phthalate-injected oocytes was cultured in IVM medium supplemented with 200 μM N-acetyl-L-cysteine (NAC), followed by transfer to M16 medium containing the same concentration of NAC. The dose of NAC was selected based on our previously established protocols [45]. All other culture conditions, including incubation durations and environmental parameters, were maintained identically across all groups.

### Immunofluorescence staining

Oocytes were fixed in 4% paraformaldehyde for 30 min at room temperature, washed three times with 1 mg/ml polyvinyl pyrrolidone (PVP; Sigma, USA) in PBS, and permeabilized in 0.1% Triton X-100 in PBS for 15 min at room temperature. After three additional washes, oocytes were blocked in 1% bovine serum albumin (BSA; Sigma-Aldrich, USA) for 1 h at room temperature. For α-tubulin and F-actin staining, oocytes were incubated overnight at 4 °C with FITC-conjugated mouse anti-α-tubulin antibody (1:200; Sigma, USA). After three washes with PBS, they were stained with Phalloidin-Alexa Fluor 594 (Invitrogen, USA) according to the manufacturer’s instructions. Briefly, oocytes were incubated with 1× Phalloidin-Alexa Fluor 594 for 30 min at 37 °C, followed by three PBS washes. DNA was counterstained with 1 µg/mL DAPI for 5 min at room temperature. For γH2AX staining, oocytes were incubated overnight at 4 °C with anti-γH2AX antibody (1:200; Millipore, USA). After three washes with PBS, the oocytes were incubated with Alexa Fluor 488-conjugated secondary antibody (1:500; Thermo Fisher Scientific, USA) for 1 h at room temperature, and then washed three times with PBS. DNA was counterstained with 1 µg/mL DAPI for 5 min at room temperature. Finally, oocytes were washed three times and imaged using a Nikon A1R confocal microscope (Nikon Instruments Inc., NY, USA). Image acquisition and processing were performed using NIS-Elements software (Nikon).

### Measurement of reactive oxygen species (ROS) levels

Intracellular reactive oxygen species (ROS) levels in live oocytes were evaluated using the oxidation-sensitive fluorescent probe 2′,7′-dichlorofluorescin diacetate (DCFH-DA; Sigma-Aldrich, USA). Oocytes were incubated in M2 medium containing 10 µM DCFH-DA at 37 °C in a 5% COL incubator for 30 min, followed by three washes in fresh M2 medium to remove residual dye. Fluorescence was immediately detected using a Nikon A1R confocal microscope (Nikon Instruments Inc., NY, USA), and signal intensity was quantified within the region of interest (ROI) using ImageJ software (National Institutes of Health, USA).

### Measurement of mitochondrial membrane potential

Mitochondrial membrane potential in oocytes was measured using a Mitochondrial Membrane Potential Detection Kit (Biotium, Inc., CA, USA) according to the manufacturer’s instructions. Briefly, oocytes were incubated in 1× JC-1 working solution at 37 °C in a 5% COL incubator for 30 min in the dark, followed by three washes with M2 medium. Fluorescence was immediately detected using a Nikon A1R confocal microscope (Nikon Instruments Inc., NY, USA). The ratio of red to green fluorescence intensity was calculated to assess mitochondrial membrane potential, and fluorescence quantification was performed using ImageJ software (National Institutes of Health, USA).

### Smart-seq3 Library Preparation and Sequencing

Single-cell RNA sequencing was conducted using the Smart-seq3 workflow [46] to profile the transcriptomes of germinal vesicle (GV) stage oocytes from water-injected (control) and phthalate-injected groups. Each group included three independent biological replicates. Oocytes were transferred into lysis buffer supplemented with RNase inhibitor to preserve RNA integrity. Reverse transcription was initiated using an oligo(dT) primer containing a 5′ anchor sequence, and a template-switching oligonucleotide (TSO) enabled generation of full-length cDNA. The synthesized cDNA was subsequently amplified by PCR and purified with magnetic beads. The concentration of amplified cDNA was measured using the Qubit dsDNA HS Assay Kit (Thermo Fisher Scientific), and size distribution of fragments was determined using an Agilent 4200 TapeStation system. Libraries were constructed using a Nextera XT DNA Library Preparation Kit (Illumina) according to the manufacturer’s guidelines and sequenced on the Illumina MiSeq platform with 150 bp paired-end (PE150) setting. For Smart-seq3 data analysis, sequencing adapters and low-quality bases were removed using Cutadapt (v2.8) and Trimmomatic (v0.39). Cleaned reads were aligned to the mouse reference genome (mm10) via Hisat2 (v2.2.1). Differential expression analysis was conducted using DESeq2 (v1.38.3). Genes with p-value < 0.05 and |logL fold change| > 1 were considered significantly differentially expressed. Gene Ontology (GO) and KEGG pathway analysis were performed using the ClusterProfiler (v4.6.2).

### Statistical Analysis

All experiments were independently repeated at least three times. Statistical analyses were performed using GraphPad Prism 10.0 software (GraphPad Software, San Diego, CA, USA). Data in bar plots are presented as the mean ± standard error of the mean (SEM). Box plots show the median, first and third quartiles, with whiskers indicating the minimum and maximum values, and violin plots illustrate the data distribution with the median indicated. Statistical comparisons between the control and phthalate-treated groups were conducted using an unpaired two-tailed Student’s *t*-test. Statistical significance was defined as* *p* < 0.05, \*\**p* < 0.01, and *** *p* < 0.001.

## Results

### Phthalate metabolite mixture inhibits oocyte meiotic maturation

To examine the effects of phthalate metabolite mixtures on oocyte quality, GV-stage oocytes were cultured in vitro with DMSO (control), 0.4 μM and 1.4 μM phthalate metabolite mixture for 18 h. Although the rate of first polar body extrusion did not differ significantly among the three groups (Fig. S1A, B), both 0.4 μM and 1.4 μM treatments caused a significant increase in abnormal MII oocytes, characterized by chromosome misalignment, compared with the control group (Fig. S1C, D). To assess the impact of physiological relevant concentrations, fully grown oocytes within cumulus-oocyte-complexes were microinjected with either water (control) or 40 nM phthalate metabolite mixture. Most phthalate-exposed oocytes failed to undergo germinal vesicle breakdown (GVBD) after 4 hours of culture (Fig. 1B). After 18 h hours, the majority of oocytes did not extrude first polar body (Fig. 1A, and C), and a substantial fraction of the oocytes was nonviable (Fig. 1A, and D). Consistent with these findings, α-tubulin immunostaining showed that most phthalate-injected oocytes were arrested at the metaphase I (MI) stage (Fig. 1E). Together, these results demonstrate that exposure to phthalate metabolite mixture severely impairs meiotic progression.

**Figure 1.**
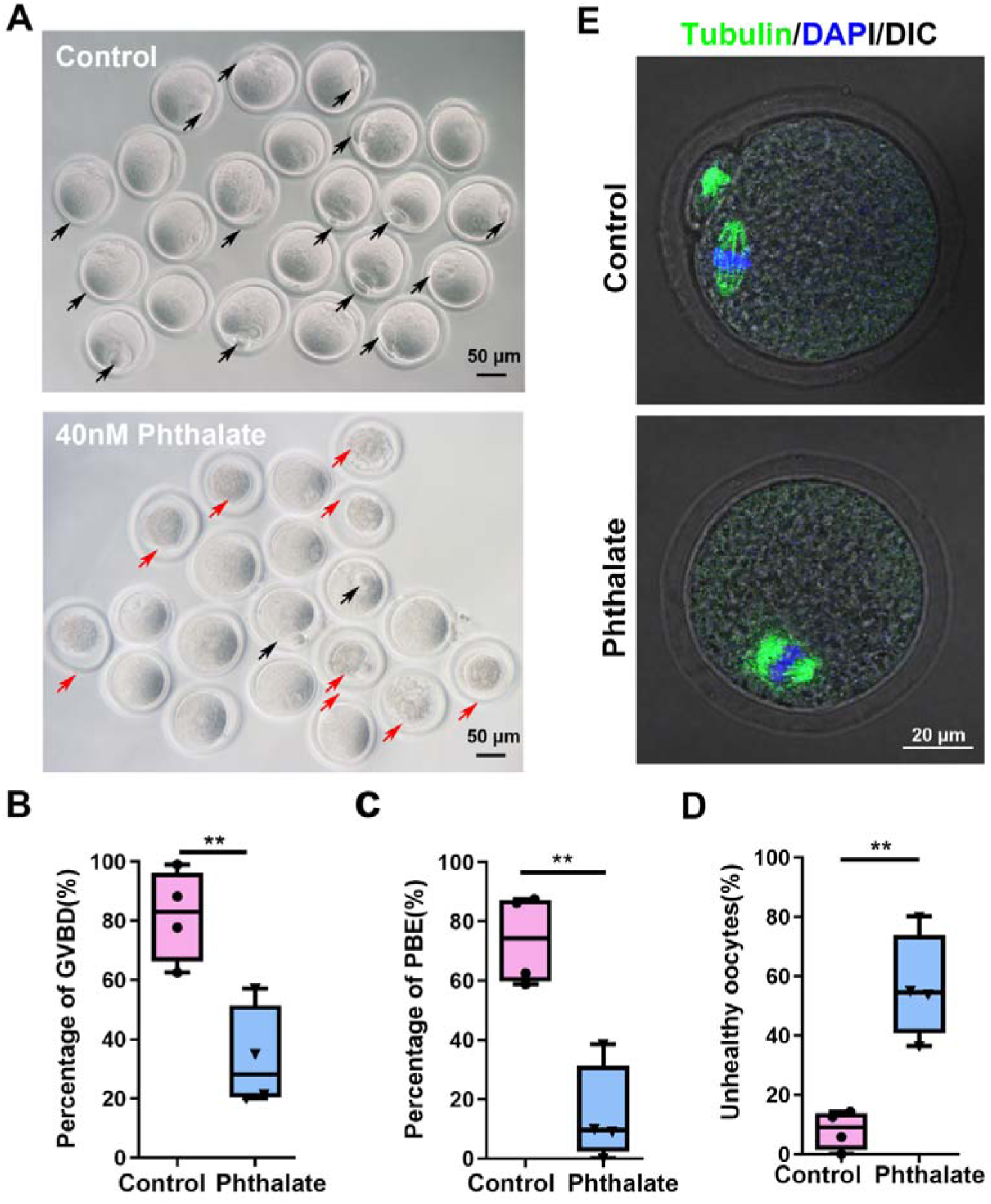
Effects of phthalate metabolite mixtures on oocyte development. (A) Representative morphology of oocytes after 18 h of in vitro culture in the control and phthalate-injected groups. Black arrows indicate MII oocytes, and red arrows indicate degenerated oocytes. Scale bar, 50 μm. (B) Quantitative analysis of GVBD rate in control and phthalate-injected groups after 4 hours of culture. Data in Boxplots show the median, first and third quartiles, and whiskers indicate the range, n=4. \*\**p* < 0.01 by two-tailed Student’s t-test. (C) Quantitative analysis of oocytes with first polar body extrusion (PBE) rate in control and phthalate-injected groups after 18 hours of culture. Boxplot representation as in (B), n=4. \*\**p* < 0.01 by two-tailed Student’s t-test. (D) Proportion of unhealthy oocytes in the control and phthalate-injected groups after 18 hours of culture. Unhealthy oocytes were defined as those exhibiting granular cytoplasm, cytoplasmic shrinkage, or detachment of the oocyte membrane from the zona pellucida (increased perivitelline space). Boxplot representation as in (B), n=4. \*\**p* < 0.01 by two-tailed Student’s t-test. (E) Immunofluorescence staining of tubulin (green) in control and phthalate-injected groups after 18 hours of culture. Nuclei are counterstained with DAPI (blue). Scale bar: 20 μm.

### Phthalate metabolite mixture injection compromises spindle integrity and cytoskeletal organization in MII oocytes

Following phthalate injection, the vast majority of oocytes underwent degeneration, and only a small proportion successfully reached the metaphase II (MII) stage. To assess the cytological quality of these surviving oocytes, we performed immunofluorescence analysis of α-tubulin on oocytes after 18 hours of *in vitro* culture. Marked spindle abnormalities were frequently observed in the phthalate-injected group (Fig. 2A), indicating severe disruption of meiotic spindle assembly. Given these defects, we next investigated whether phthalate exposure impaired the organization of the actin cytoskeleton. F-actin–a key component of the cortical cytoskeleton that regulates spindle positioning, asymmetric division, and cytoplasmic organization–was visualized using Phalloidin staining. The majority of control MII oocytes exhibited a well-defined actin cap. In contrast, most MII phthalate-injected oocytes displayed abnormal F-actin caps that displayed reduced fluorescence intensity (Fig. 2B, C). Quantitative analysis showed a significant increase in the proportion of oocytes with abnormal F-actin caps in the phthalate-injected group compared with the control group (Fig. 2D). Collectively, these findings indicate that, although a few oocytes can progress to the MII stage following phthalate exposure, they exhibit pronounced spindle abnormalities and cytoskeletal disorganization, which may underlie their compromised developmental competence.

**Figure 2.**
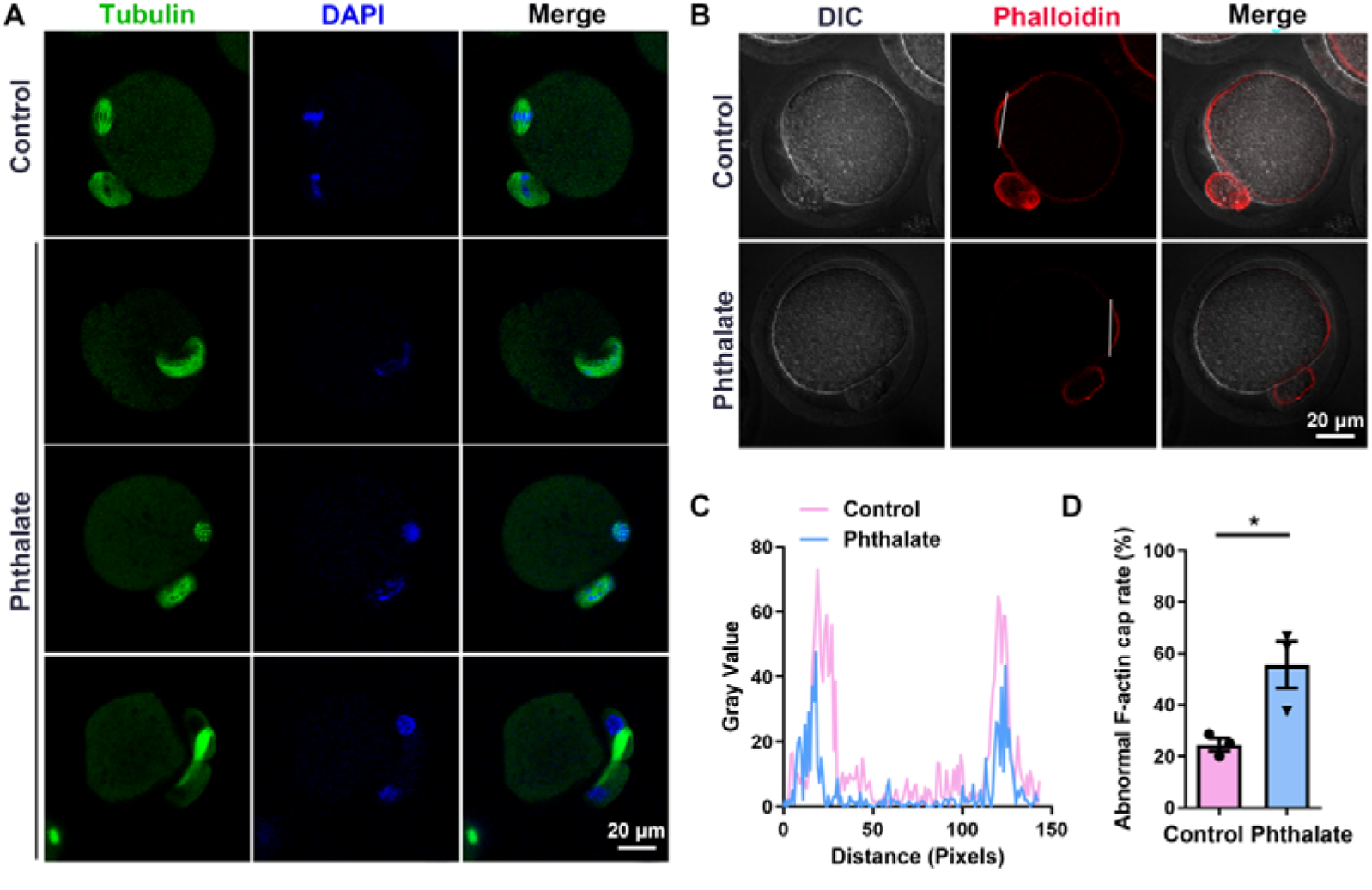
Phthalate exposure induces spindle abnormalities and disrupts cortical F-actin organization in MII oocytes. (A) Representative immunofluorescence images showing spindle morphology in MII oocytes from the water-injected (control) and phthalate-injected groups. Oocytes were stained with α-tubulin (green) to visualize spindle structures, and DNA was counterstained with DAPI (blue). Abnormal spindle morphology was frequently observed in the phthalate-injected group. Scale bar: 20 μm. (B) Fluorescence images of F-actin in control and phthalate-injected oocytes after 18 hours culture. Phalloidin (red) marks F-actin. White lines indicate cap regions used for fluorescence measurements. Scale bar, 20 μm. (C) F-actin fluorescence intensity profiles in control and phthalate-injected groups, measured along the white lines shown in (B) (D) The ratio of abnormal F-actin cap structures after 18 hours of in vitro culture in control and phthalate-injected groups. Data were presented as mean ± SEM from three independent experiments (n=3). \**p* < 0.05 by two-tailed Student’s t-test.

### Phthalate metabolite mixture disrupts spindle morphology and chromosome alignment in MI oocytes

Given that most oocytes in the phthalate metabolite mixture-exposed group remained arrested at the metaphase I (MI) stage after 18 hours of culture (Fig. 1E), we next examined MI-stage oocytes to gain mechanistic insight into phthalate-induced meiotic failure. Immunofluorescence staining was performed after 8 hours of culture to assess spindle morphology. At this time point, a considerable proportion of oocytes exhibited signs of degeneration (Fig. 3A), only the remaining viable oocytes were included in the subsequent immunostaining analyses. In the control oocytes, the meiotic spindle displayed the expected barrel-shaped structure with chromosomes well aligned along the equatorial plate (Fig. 3B). In contrast, oocytes exposed to phthalate metabolite mixture displayed pronounced spindle abnormalities accompanied by chromosome misalignment (Fig. 3B). Quantitative analysis further revealed a significant reduction in spindle length in the phthalate-exposed oocytes (Fig. 3C, D), whereas spindle width remained unaltered (Fig. 3C, and E) compared to controls. Consistently, the MI plate width was markedly increased following phthalate exposure compared to control (Fig. 3C, and F). Collectively, these findings demonstrate that phthalate metabolite mixture severely disrupts meiotic spindle assembly and compromises chromosome alignment, providing a mechanistic basis for the observed failures in meiotic progression.

**Figure 3.**
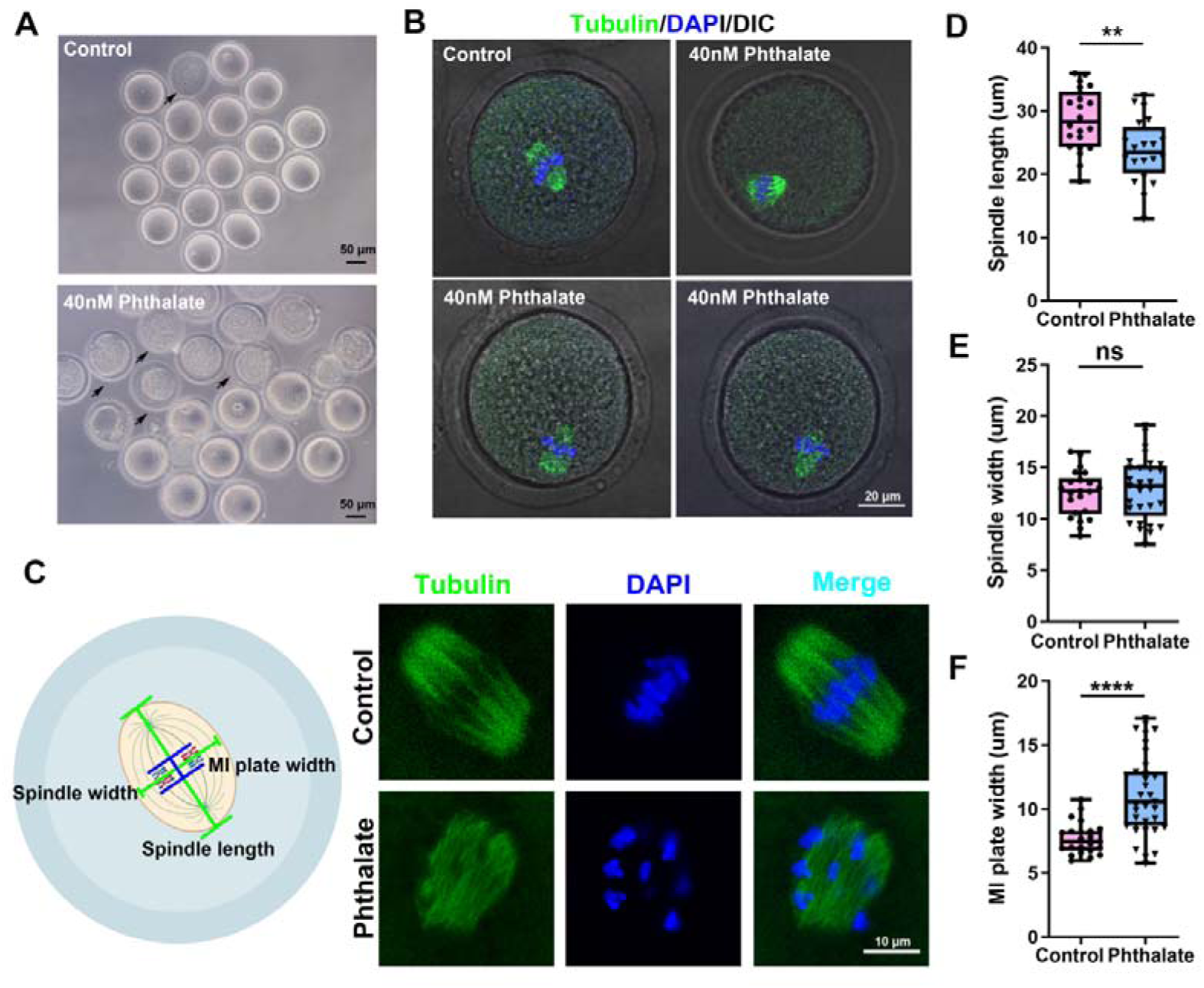
Phthalate exposure impairs meiotic spindle assembly and chromosome alignment in MI oocytes. (A) Representative brightfield images of oocytes after 8 hours of *in vitro* culture in control and phthalate-injected groups. Arrows indicate degenerated oocytes. Scale bar, 50 μm. (B) Immunofluorescence staining of α-tubulin (green) in control and phthalate-injected groups after 8 hours of culture. DNA were counterstained with DAPI (blue). Scale bar: 20 μm. (C) Schematic illustration showing the measurement of spindle length, spindle width, and MI plate width (left). Spindle length was measured as the distance between the two spindle poles, while spindle width as the microtubule width at the MI plate. MI plate width was defined as the distance between the two outer edges of the DNA/chromosomes. Representative images of metaphase I (MI) spindle and chromosome morphology in control and phthalate-injected oocytes (right). Spindles were stained with α-tubulin (green), and DNA/chromosomes were counterstained with DAPI (blue). Scale bar: 10 μm. (D) Quantitative analysis of MI spindle length in control and phthalate-injected groups. Data in Boxplots show the median, first and third quartiles, and whiskers indicate the range, n=22 for control and 20 for phthalate-injected groups from three biological replicates. \*\**p* < 0.01 by two-tailed Student’s t-test. (E) Quantitative analysis of meiotic spindle width in control and phthalate-injected groups. Data in boxplots show the median, first and third quartiles, and whiskers indicate the range, n=22 for control and 29 for phthalate-injected groups from three biological replicates. ns, not significant (p ≥ 0.05) by two-tailed Student’s t-test. (F) Quantitative analysis of MI plate width in in control and phthalate-injected groups. Data in boxplots show the median, first and third quartiles, and whiskers indicate the range, n=23 for control and 35 for phthalate-injected groups from three biological replicates. \*\*\*\**p* < 0.0001 by two-tailed Student’s t-test.

### Phthalate metabolite mixture disrupts actin cytoskeleton in MI oocytes

Given the critical interplay between microtubules and actin filaments during meiotic progression, we next examined whether exposure to phthalate metabolite mixture perturbs actin cytoskeleton organization in MI oocytes. F-actin was visualized using phalloidin staining to assess whether altered actin dynamics contribute to the phthalate-induced meiotic defects. In control oocytes, robust F-actin signals were prominently enriched along the plasma membrane, forming a well-defined cortical actin layer, as illustrated by the fluorescence intensity profiles across the oocyte (Fig. 4A, B). This cortical network is essential for maintaining oocyte shape, facilitating spindle positioning, and supporting polar body extrusion. In contrast, MI oocytes exposed to phthalate metabolite mixture displayed a markedly reduced F-actin signals (Fig. 4A, B). Quantitative analysis further confirmed significantly decreased F-actin fluorescence intensity in phthalate-injected oocytes compared with controls (Fig. 4C). Collectively, these results demonstrate that phthalate metabolite mixture compromises cortical actin formation during oocyte meiosis, highlighting a cytoskeletal mechanism that likely contributes to phthalate-induced meiotic defects and impaired oocyte quality.

**Figure 4.**
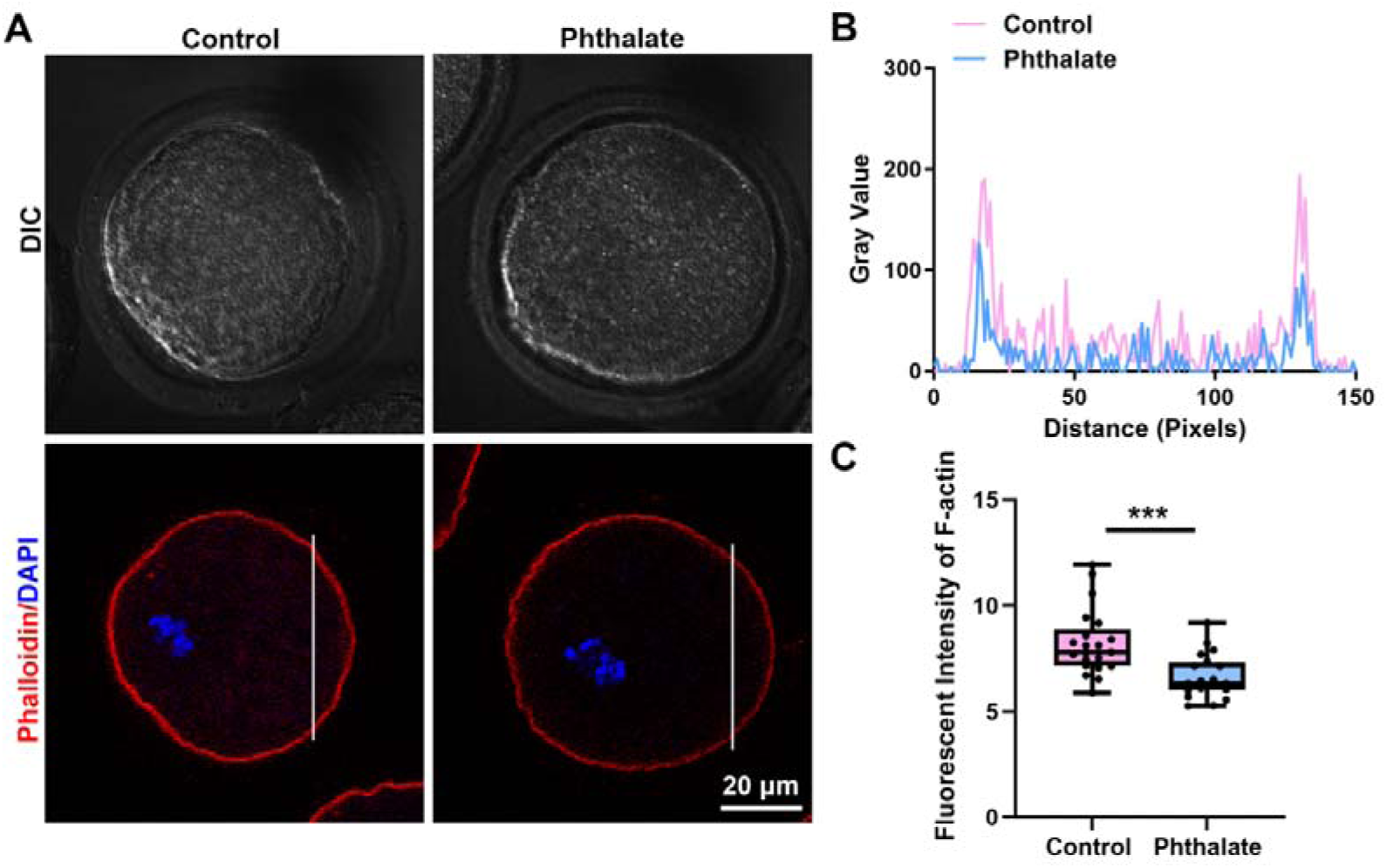
Effects of phthalate metabolite mixtures on actin cytoskeleton formation. (A) Representative fluorescence images of F-actin in control and phthalate-injected oocytes after 8 hours of *in vitro* culture. Phalloidin (red) marks F-actin. Scale bar, 20 μm. (B) Fluorescence intensity profiles of F-actin in control and phthalate-injected oocytes, measured along the white lines shown in (A). (C) Quantification of F-actin fluorescence intensity at the plasma membrane in control and phthalate-injected oocytes. Data in boxplots show the median, first and third quartiles, and whiskers indicate the range, n=21 for control and 20 for phthalate-injected groups from three biological replicates. \*\*\**p* < 0.001 by two-tailed Student’s t-test.

### Phthalate metabolite mixture affects DNA damage repair in oocytes

Previous studies have shown that DEHP exposure can impair DNA damage repair in fetal mouse oocytes [47], raising the possibility that phthalate compounds may compromise genomic integrity. Thus, we hypothesized that exposure to phthalate metabolite mixture similarly affects DNA damage repair in MI oocytes. To test this hypothesis, oocytes were cultured for 8 hours post-injection and subsequently subjected to γ-H2AX staining, a well-established marker of DNA double-strand breaks (DSBs). Imaging analysis revealed that the γ-H2AX signal was markedly enhanced in phthalate-exposed oocytes compared with controls (Fig. S2A). Quantitative analysis further confirmed a significant elevation in γ-H2AX fluorescence intensity in the phthalate-injected group compared to control (Fig. S2B), indicating that phthalate metabolite mixture affects DNA damage repair during oocyte maturation. Together, these results suggest that phthalate metabolite mixtures can affects DNA damage repair in oocytes, potentially contributing to meiotic arrest and reduced oocyte quality.

### Phthalate metabolite mixture impairs mitochondrial function in oocyte

In the phthalate-injected group, the GVBD rate was significantly reduced compared to controls, suggesting that exposure to phthalate metabolite mixture may hinder GV oocyte maturation. Given that mitochondrial dysfunction is common consequence of toxicant exposure and that the phthalate DiNP has been shown to alter mitochondrial dynamics in porcine oocytes [48], we next investigated whether phthalate metabolite mixtures similarly affect mitochondrial function in mouse oocytes. After 2 hours of *in vitro* culture, oocytes were collected for JC-1 staining to evaluate the mitochondrial membrane potential (ΔΨm), an indicator of mitochondrial function. JC-1 emits green fluorescence in its monomeric form and red fluorescence when it forms J-aggregates in mitochondria with high membrane potential. Therefore, the red-to-green fluorescence ratio reflects mitochondrial polarization status. As shown in Fig. 5A, JC-1 fluorescence patterns differed markedly between control and phthalate-exposed oocytes. Quantitative analysis revealed a significant decrease in the red/green fluorescence ratio in the phthalate-injected group compared with controls (Fig. 5B). These results demonstrate that phthalate metabolite mixture compromises mitochondrial function in mouse oocytes.

**Figure 5.**
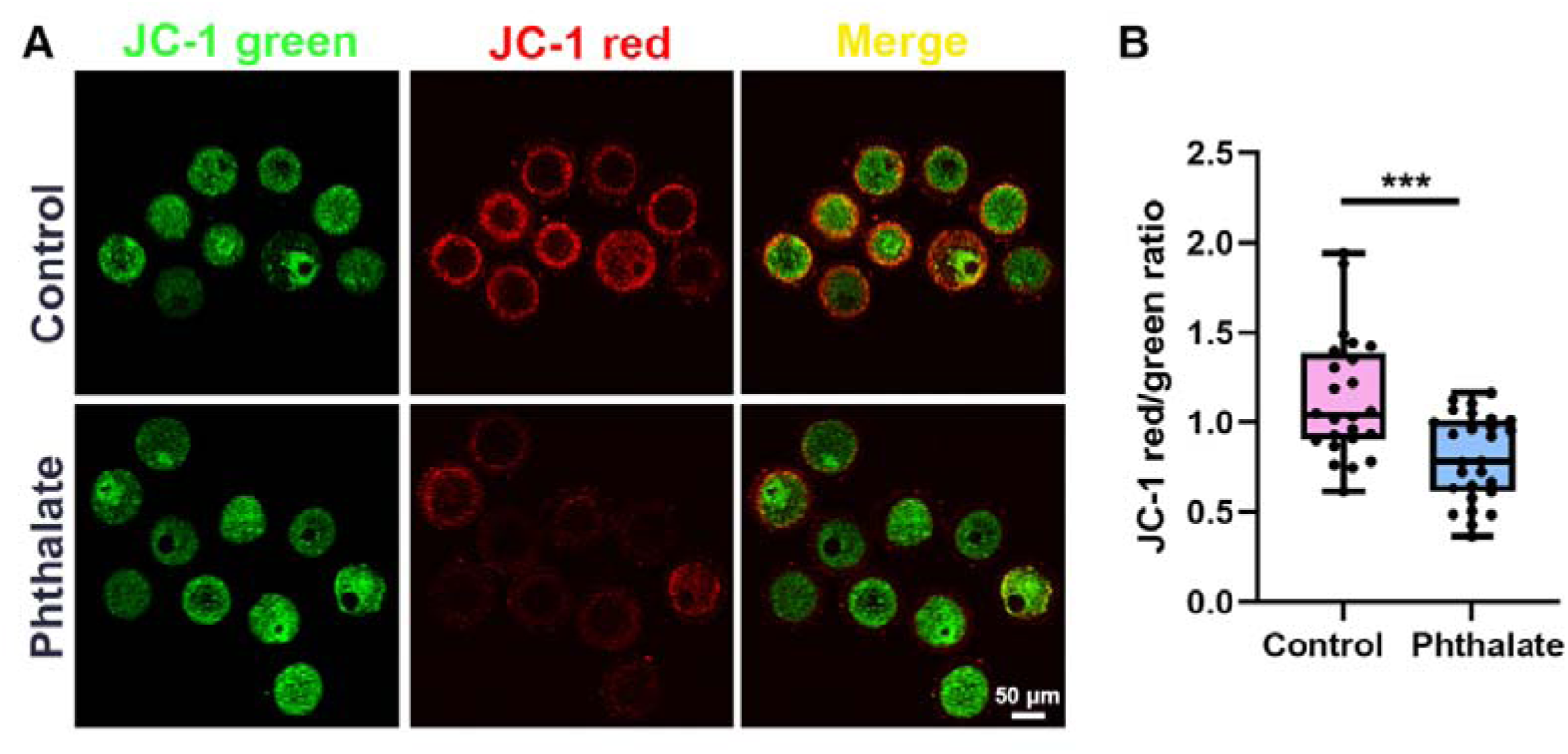
Phthalate metabolite mixture affects mitochondrial membrane potential. (A) Representative JC-1 staining images showing mitochondrial membrane potential in control and phthalate-injected oocytes after 2-hour culture. Green fluorescence corresponds to JC-1 monomers (low membrane potential); while red fluorescence indicates JC-1 aggregates (high membrane potential). Scale bar: 50 µm. (B) Quantification of mitochondrial membrane potential is represented by the JC-1 red/green fluorescence ratio (n = 24 for control and 27 for phthalate-injected oocytes from three biological replicates, Data in Boxplots show the median, first and third quartiles, and whiskers indicate the range, *** *p* <0.001).

### Phthalate metabolite mixture increases ROS production and induces DNA damage in oocytes

Previous studies have reported that phthalates can induce oxidative stress [48, 49]. Given that the phthalate metabolite mixture disrupts mitochondrial function in our study, we hypothesized that such mitochondrial disruption would consequently elevate ROS levels in oocytes. To test this, we measured ROS levels using DCFH-DA, a well-established fluorescent probe for ROS detection. DCFH-DA is a non-fluorescent, cell-permeable probe that readily enters live cells. Intracellular esterases remove the diacetate groups, converting DCFH-DA into DCFH, which is trapped inside the cell but remains non-fluorescent. Reactive oxygen species then oxidize DCFH to form DCF, a highly fluorescent compound. Thus, the accumulation of DCF fluorescence reflects intracellular oxidative activity. Our results showed that DCF fluorescence was markedly stronger in phthalate-injected oocytes compared with controls (Fig. 6A). Quantitative analysis confirmed a significant increase in fluorescent intensity in the treated group (Fig. 6B), indicating elevated ROS levels following exposure. Because excessive ROS can lead to DNA damage, we next examined whether phthalate metabolite mixture exposure induced DNA damage in oocytes. GV-stage oocytes were harvested for γ-H2AX immunostaining, which marks DNA double-strand breaks. The γ-H2AX signal was markedly stronger in phthalate-injected oocytes compared with controls (Fig. 6C). Quantification confirmed a significant increase in average γ-H2AX intensity in phthalate exposed oocytes compared to controls (Fig. 6D). Together, these results indicate that exposure to the phthalate metabolite mixture elevates ROS levels and induces DNA damage in mouse oocytes.

**Figure 6.**
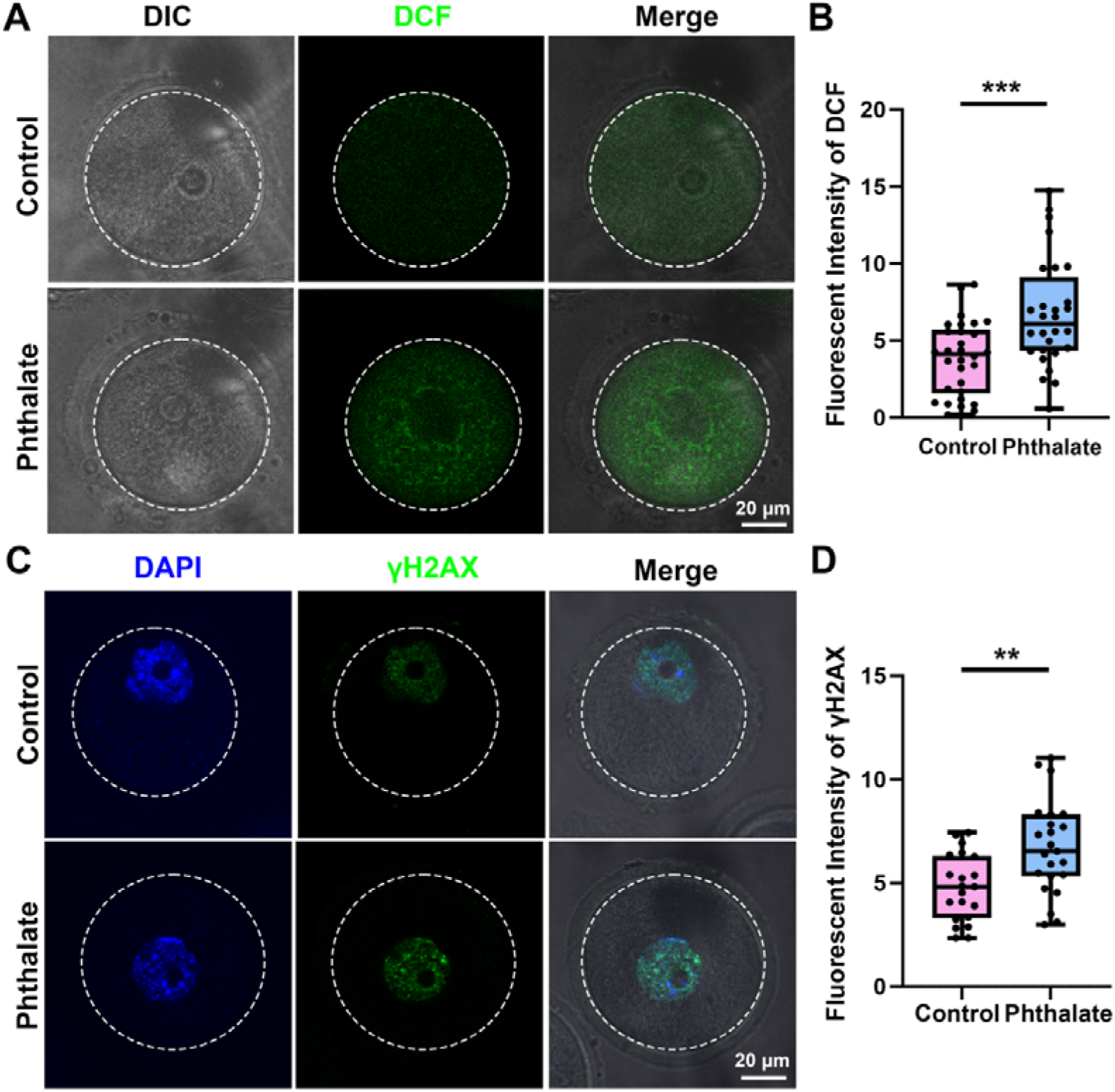
Exposure to phthalate metabolite mixture induces oxidative stress and DNA damage in mouse oocytes. (A) Representative images of DCF fluorescence (green) in control and phthalate metabolite mixture-exposed oocytes. DIC, differential interference contrast. Scale bar = 20 μm. (B) Quantification of DCF fluorescence intensity in control and phthalate metabolite mixture–treated oocytes. ROS levels were analyzed in 30 control oocytes and 28 treated oocytes. Data are represented as mean ± SEM of at least three independent experiments. t-test, \*\*\**P* < 0.001, compared with control. (C) Representative images showing γ-H2AX (green) and DNA (blue) in control and phthalate injected oocytes. Scale bar = 20 μm. (D) Quantitative analysis of average γ-H2AX (green) intensity in control and phthalate metabolite mixture-exposed oocytes. A total of 21 control oocytes and 23 treated oocytes were analyzed. Data are represented as mean ± SEM of at least three independent experiments. t-test, \*\**P* < 0.01, compared with control.

### Smart RNA-seq reveals transcriptomic alterations in oocytes after phthalate metabolite mixture exposure

To elucidate the molecular mechanisms underlying the effects of phthalate-induced defects in oocyte meiotic maturation, Smart RNA sequencing (Smart RNA-seq3) was performed to compare transcriptomic profiles between control and phthalate-exposed oocytes. Principal component analysis (PCA) revealed a clear separation between the two groups, indicating substantial transcriptomic reprogramming induced by phthalate exposure (Fig. 7A). A total of 201 differentially expressed genes (DEGs) were identified, including 117 downregulated and 84 upregulated genes, as shown in the volcano plot and heatmap analyses (Fig. 7B, C). Consistent with our earlier observations of mitochondrial dysfunction, cytoskeletal disorganization, and DNA damage, many DEGs were associated with mitochondrial activity, cytoskeletal organization, and DNA repair pathways (Fig. 7D), suggesting that phthalate exposure perturbs these cellular processes at the transcriptional level.

**Figure 7.**
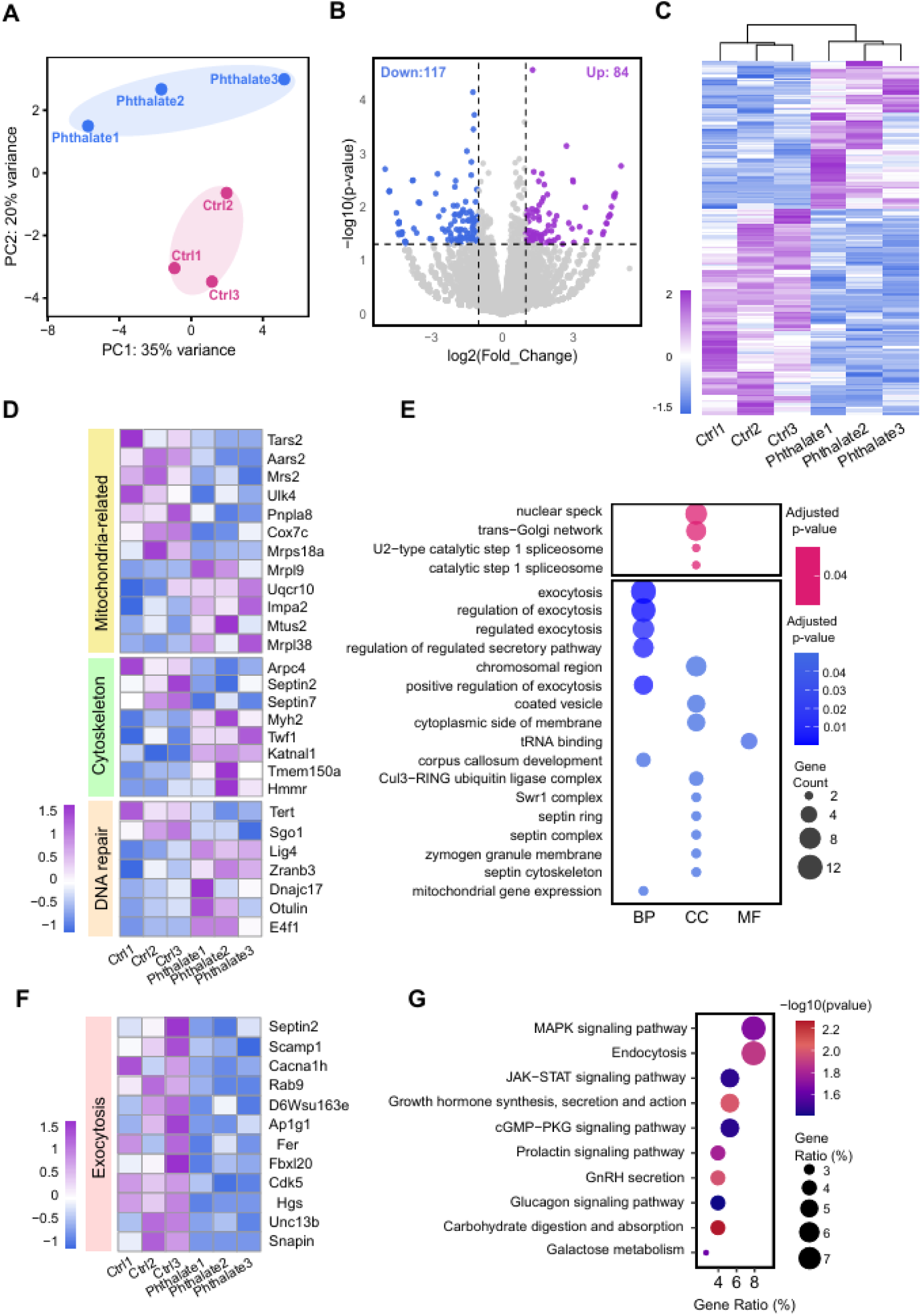
Transcriptomic analysis of oocytes exposed to phthalate metabolite mixture. (A) Principal component analysis (PCA) of phthalate-injected GV oocytes (n = 3) and water-injected controls (n = 3), showing clear separation between the two groups. (B) Volcano plot showing differentially expressed genes (DEGs) between phthalate-injected and control oocytes. Purple dots indicate upregulated genes, and blue dots indicate downregulated genes. (C) Heatmap illustrating the expression levels of DEGs across samples. Each column represents an individual sample, and each row corresponds to a DEG. Color gradient from light blue to light purple reflect increasing expression levels. (D) Representative DEGs associated with mitochondrial function, cytoskeletal organization, and DNA repair. (E) Gene Ontology (GO) enrichment analysis of DEGs, showing significantly enriched terms in biological processes (BP), cellular components (CC), and molecular functions (MF). (F) Heatmap displaying DEGs involved in the exocytosis regulation. (G) Selected enriched KEGG pathways associated with the DEGs.

To further explore the biological implications of these DEGs, Gene Ontology (GO) and Kyoto Encyclopedia of Genes and Genomes (KEGG) pathway enrichment analyses were conducted. GO analysis revealed significant enrichment of DEGs in biological processes related to exocytosis, regulation of the secretory pathway, cytoskeletal organization, and mitochondrial gene expression. Mitochondria-related DEGs such as calcium-independent phospholipase A2-gamma (*Pnpla8*) has been reported to have an antioxidant role in limiting mitochondrial superoxide production, suggesting potential alterations in redox regulation after phthalate exposure [50]. Enriched cellular component terms included the trans-Golgi network, coated vesicles, and the septin cytoskeleton (Fig. 7E). Notably, several exocytosis-related genes, including secretory carrier-associated membrane protein 1 (*Scamp1)*, *Snapin*, *Unc13b*, and *Rab9*, exhibited altered expression following phthalate exposure (Fig. 7F). KEGG pathway analysis further demonstrated that DEGs were primarily enriched in the MAPK signaling pathway, JAK-STAT signaling pathway, cGMP-PKG signaling pathway, and GnRH secretion pathway (Fig. 7G), which are closely associated with cell signaling, follicular development, oocyte maturation, and hormone regulation. Together, these findings provide transcriptomic evidence that phthalate exposure disrupts key molecular pathways critical for oocyte developmental competence and reproductive function.

### N-acetyl-L-cysteine rescues meiotic maturation defects induced by phthalate metabolite mixture exposure

Transcriptomic analysis revealed significant alterations in the mitochondria-related pathway in phthalate-metabolite-mixture-exposed oocytes (Fig. 7D). Consistent with these findings, our fluorescence staining assays demonstrated elevated intracellular ROS levels (Fig. 6 A, B), reduced mitochondrial membrane potential (Fig. 5), and increased DNA damage (Fig. 6 C, D) following phthalate exposure, suggesting disrupted redox homeostasis and oxidative stress-associated cellular dysfunction. To further evaluate the functional contribution of oxidative stress in phthalate-induced meiotic impairment and to explore potential intervention strategies, we investigated whether antioxidant supplementation could alleviate the impaired developmental competence of exposed oocytes. N-acetyl-L-cysteine (NAC), a well-characterized ROS scavenger and glutathione precursor, was employed to reduce intracellular oxidative stress [45]. Oocytes were treated with the phthalate metabolite mixture in the presence or absence of NAC (200 μM), and meiotic progression was assessed by monitoring GVBD and PBE.

As previously observed, phthalate exposure significantly suppressed both GVBD and PBE rates compared to controls. Notably, NAC supplementation partially alleviated these inhibitory effects, resulting in improved GVBD and PBE rates in the co-treatment group (Fig. 8A–B). Quantitative analysis further demonstrated that the PBE rate was significantly increased in the NAC-treated group compared with the phthalate-only group and that it approached control levels (Fig. 8C). Together, these findings provide functional evidence supporting the involvement of oxidative stress in phthalate-induced meiotic impairment and suggest that reducing ROS accumulation can partially restore oocyte maturation capacity in phthalate exposed oocytes.

**Figure 8.**
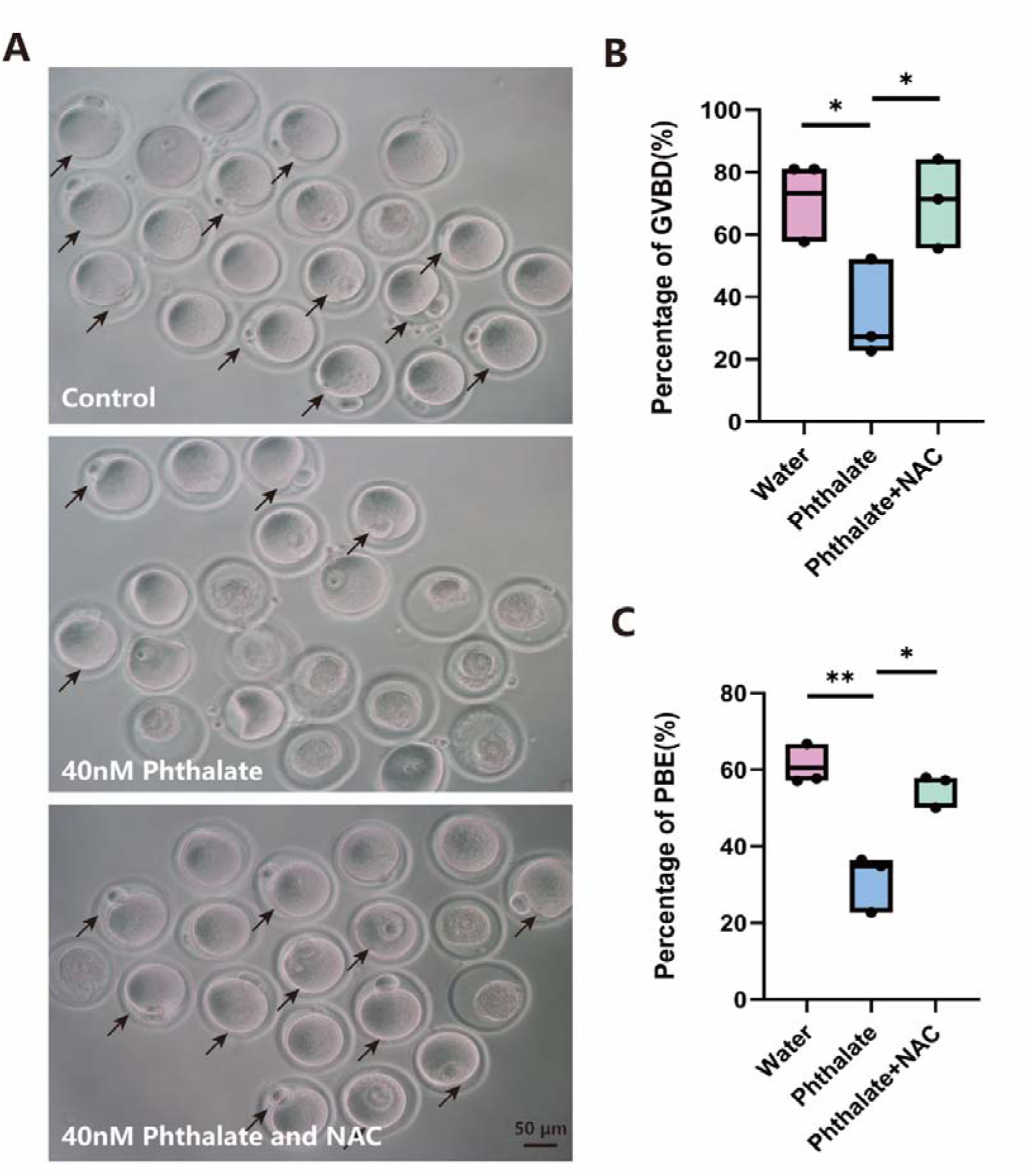
NAC rescues meiotic maturation defects induced by phthalate metabolite mixture exposure. (A) Representative morphology of oocytes after 18 h of *in vitro* culture in the control, phthalate-treated, and phthalate + NAC groups. Black arrows indicate MII oocytes. Scale bar, 50 μm. (B) Quantitative analysis of GVBD rate in the control, phthalate-treated, and phthalate + NAC groups after 4 hours of culture. Data are presented as individual biological replicates with mean values indicated, n = 3. *p < 0.05, **p < 0.01 by two-tailed Student’s t-test. (C) Quantitative analysis of PBE rates in control, phthalate-treated, and phthalate + NAC groups after 18 hours of culture. Data are presented as in individual biological replicates with mean values indicated; n = 3. *p < 0.05, **p < 0.01 by two-tailed Student’s t-test.

## Discussion

Phthalates are widely recognized reproductive toxicants that impair ovarian physiology and female fertility [51]. Here, we demonstrate that even a low-concentration phthalate metabolite mixture (40 nM; 36.7% MEP, 19.4% MEHP, 15.3% MBP, 10.2% MiBP, 10.2% MiNP, and 8.2% MBzP) markedly reduces the developmental competence of mouse oocytes, leading to decreased GVBD and failure of first polar-body extrusion. These defects are accompanied by disrupted spindle organization, compromised mitochondrial function, increased oxidative stress, and DNA damage in exposed oocytes. Transcriptomic profiling further highlighted extensive changes in molecular pathways vital for oocyte maturation.

Previous studies on phthalate reproductive toxicity have primarily focused on the effects of individual compounds. MEHP, for example, impairs *in vitro* mouse oocyte maturation and subsequent embryonic development [52], and follicular fluid containing MEHP with reduced estradiol levels led to fewer bovine oocytes reaching the MII stage during *in vitro* maturation [53]. Neonatal DEHP exposure disrupts germ cell nest breakdown and primordial follicle formation through multiple underlying mechanisms [54]. Similarly, DBP also compromises oocyte maturation, as evidenced by decreased GVBD and polar-body extrusion rates in both *in vitro* and *in vivo* mouse models [55], and gestational exposure to DBP interferes with meiotic prophase I progression in fetal mouse oocytes, particularly from the zygotene to pachytene stages [27]. In porcine oocytes, MBP induced meiotic defects and apoptosis by modulating mitochondrial-ER (endoplasmic reticulum) interactions [56]. Although these studies demonstrate the reproductive toxicity of individual phthalates, they do not reflect real-world exposure, in which humans encounter complex mixtures of phthalates through various consumer products. Indeed, mixtures of phthalate metabolites have been detected in human urine, serum, and follicular fluid [15, 32, 34, 57, 58], indicating oocytes are exposed to multiple phthalates within the ovarian microenvironment. Our study addresses this gap by directly examining the effects of a physiologically relevant phthalate mixture, revealing that combined exposure induces pronounced oocyte developmental defects. These data highlight the significance of mixture-based toxicological assessment to more accurately evaluate reproductive health risks.

From a public health perspective, our findings raise concerns about the reproductive consequences of environmental exposure to phthalate mixtures. Since phthalate metabolites have been consistently detected in human follicular fluid, oocytes are directly exposed to phthalates throughout folliculogenesis. Previous studies examining follicular fluid from women undergoing fertility treatment have reported that the concentrations of detected phthalate metabolites (including MEP, MEHP, MBuP, and MMP) were all below 15 ng/mL, with residual MEHP levels ranging from approximately 20 to 51 nM [59]. Another epidemiological study similarly identified MEHP and MBP as dominant metabolites in human follicular fluid, with median concentrations of 2.80 ng/mL (10.1 nM) and 2.05 ng/mL (9.23 nM), respectively [15]. Collectively, these previous studies indicate that phthalate metabolite concentrations in follicular fluid generally fall within the nanomolar range. However, most experimental studies investigating the reproductive toxicity of phthalates have employed concentrations several orders of magnitude higher. For example, exposure to 200 or 400 μM MEHP has been shown to impair mouse oocyte maturation in vitro [52]. Similarly, *in vitro* fetal mouse ovarian culture studies have demonstrated that exposure to 10 or 100 μM DEHP disrupts meiotic progression from the pachytene to diplotene stage [47]. Moreover, exposure to a phthalate mixture at 1-500 μg/mL was reported to inhibit follicle growth and induce oocyte fragmentation in isolated antral follicles [28]. In contrast, our study used a nanomolar-level, physiologically relevant phthalate metabolite mixture (40 nM), providing a more accurate representation of human exposure and increasing the translational relevance.

Oocyte quality is a key determinant of female fertility, and even subtle disruptions in meiotic progression may cause lasting adverse reproductive consequences. Proper spindle assembly and actin dynamics are essential for accurate chromosome alignment and segregation during meiosis. Consistent with this, previous studies have demonstrated that phthalates adversely affect cytoskeletal dynamics in oocytes. For example, DBP at 500-800 μM disrupts spindle morphology and damages the oocyte cytoskeletal structure [55]. The DBP primary metabolite, MBP, similarly induces spindle abnormalities, and alters cytoskeleton-associated pathways at concentrations as low as 50 μM [56]. Recent evidence indicates that exposure to DiNP or diisobutyl phthalate (DiBP) also impairs meiotic maturation in porcine oocytes by disrupting spindle/chromosome architecture and actin organization [48, 60]. In line with these findings, our data show that even nanomolar concentrations of a phthalate metabolite mixture are sufficient to disrupt spindle morphology and actin distribution. These results suggest that low-level environmental exposure may perturb the cytoskeletal machinery required for oocyte maturation, thereby compromising meiotic maturation and ultimately threatening female fertility.

Mitochondria are primary targets of environmental toxicants [61], and maturing oocytes contain exceptionally high mitochondrial content [62]. Numerous studies have shown that phthalates can impair mitochondrial function, leading to elevated ROS levels, which in turn damage spindle structures and DNA integrity. DiNP exposure, for instance, induces mitochondrial dysfunction and increases ROS, resulting in DNA damage in porcine oocytes [48]. Similarly, DBP elevates oxidative stress, downregulates key meiotic regulators, and promotes oocyte apoptosis [27]. MBP disrupts mitochondrial-endoplasmic reticulum interactions in porcine oocytes and triggers apoptosis [56], whereas DiBP impairs mitochondrial function and increases ROS levels [60]. Consistent with these findings, our study demonstrates that a low-concentration phthalate metabolite mixture significantly disrupts mitochondrial function, elevates ROS levels, and induces DNA damage in mouse oocytes. Together, these results suggest that phthalate metabolite mixtures target mitochondria, leading to oxidative stress, subsequently compromising spindle structures and DNA integrity, ultimately impairing oocyte development.

Beyond these localized mitochondrial and structural defects, our transcriptomic analysis revealed broader alterations in molecular pathways relevant to oocyte developmental competence. Specifically, GO analysis revealed enrichment of genes involved in exocytosis and vesicle trafficking (e.g., *Scamp1*, *Snapin*, *Unc13b*, and *Rab9*), processes that have been implicated in oocyte cytoplasmic maturation. Concurrently, KEGG analysis indicated enrichment of several signaling pathways, including MAPK, JAK–STAT, and cGMP–PKG pathways, all of which are known to participate in meiotic progression and cellular signaling. Given that vesicle trafficking depends on coordinated cytoskeletal organization, and that cellular redox homeostasis is critical for maintaining oocyte physiology, these transcriptomic alterations may be linked to the mitochondrial dysfunction and cytoskeletal disruption observed in this study. To further explore whether phthalate-mixture treatment induces oxidative stress, we performed a functional rescue experiment using the antioxidant NAC. NAC treatment partially alleviated phthalate-induced meiotic defects, supporting redox imbalance in the phthalate-treated oocyte dysfunctions. Although NAC rescue does not establish a direct causal relationship between oxidative stress and individual transcriptomic alterations, these findings provide functional evidence supporting the biological relevance of the oxidative stress-associated dysfunction identified through our transcriptomic and phenotypic analyses.

Despite the strengths of our study, several limitations should be acknowledged. First, although RNA-seq analysis identified multiple affected pathways, our experimental validation primarily focused on mitochondrial dysfunction and oxidative stress, as these pathways were strongly supported by both phenotypic observations (Fig. 5 and 6) and existing literature. Other enriched pathways, such as exocytosis, vesicle trafficking, and specific signal transduction cascades, were not directly validated, as many of the identified candidate genes remain poorly characterized in oocytes. In particular, their expression patterns, subcellular localization, and functional roles in oocyte biology are not well defined. Addressing these questions would require substantial preliminary characterization and dedicated experimental approaches, which were beyond the scope of the present work. Second, although the microinjection approach allows precise control of intracellular phthalate concentrations, it does not fully recapitulate chronic, low-dose *in vivo* exposure or metabolic processing of phthalates. Thus, future studies incorporating *in vivo* exposure models, alternative mixture designs, and human cohort data will be needed to refine dose-response relationships and improve translational relevance.

In conclusion, our findings demonstrate that exposure to a physiologically relevant phthalate metabolite mixture impairs oocyte maturation by disrupting cytoskeletal integrity, mitochondrial function, and key signaling pathways. These findings advance our understanding of how environmental relevant phthalate mixtures compromise female gamete quality and underscore the urgent need for mixture-based risk assessment frameworks to better protect women’s reproductive health.

## Author contributions

Juan Dong and Huanyu Qiao conceived and designed the study. Juan Dong, Vidhi Patel, Hasanur Alam, and Wenjie Yang performed all bench experiments. Juan Dong, Shuangqi Wang, and Leyi Wang collected and analyzed the data. Juan Dong, Anika Roy and Huanyu Qiao wrote the paper. Juan Dong and Huanyu Qiao supervised the project. All authors read and approved the final manuscript.

## Acknowledgements

This work was supported by NIH R00 HD082375, NIH R01 GM135549, NIH R35ES034988, and CCIL Seed Grant.

## Ethics

All animal experiments were approved by University of Illinois Urbana-Champaign (UIUC) Institutional Animal Care and Use Committee.

## Data availability

All data supporting this study are available within the article and its supplementary information. Single-cell RNA-seq data are deposited in the GEO (Gene Expression Omnibus) database with the accession number GSE315485. Additional datasets or related information are available from the corresponding authors upon reasonable request.

## References

1. Organization, W.H., Infertility prevalence estimates, 1990–2021. 2023: World Health Organization.

2. Telfer, E.E., et al., Making a good egg: human oocyte health, aging, and in vitro development. Physiol Rev, 2023. 103(4): p. 2623–2677.

3. Tan, J.H., et al., Chromatin configurations in the germinal vesicle of mammalian oocytes. Mol Hum Reprod, 2009. 15(1): p. 1–9.

4. Pan, B. and J. Li, The art of oocyte meiotic arrest regulation. Reprod Biol Endocrinol, 2019. 17(1): p. 8.

5. Li, J., W.P. Qian, and Q.Y. Sun, Cyclins regulating oocyte meiotic cell cycle progressiondagger. Biol Reprod, 2019. 101(5): p. 878–881.

6. Mogessie, B., Advances and surprises in a decade of oocyte meiosis research. Essays Biochem, 2020. 64(2): p. 263–275.

7. Sharma, A., et al., Journey of oocyte from metaphase-I to metaphase-II stage in mammals. J Cell Physiol, 2018. 233(8): p. 5530–5536.

8. Malott, K.F. and U. Luderer, Toxicant effects on mammalian oocyte mitochondriadagger. Biol Reprod, 2021. 104(4): p. 784–793.

9. Heudorf, U., V. Mersch-Sundermann, and J. Angerer, Phthalates: toxicology and exposure. Int J Hyg Environ Health, 2007. 210(5): p. 623–34.

10. Guo, Y. and K. Kannan, A survey of phthalates and parabens in personal care products from the United States and its implications for human exposure. Environ Sci Technol, 2013. 47(24): p. 14442–9.

11. Frederiksen, H., N.E. Skakkebaek, and A.M. Andersson, Metabolism of phthalates in humans. Mol Nutr Food Res, 2007. 51(7): p. 899–911.

12. Koch, H.M. and A.M. Calafat, Human body burdens of chemicals used in plastic manufacture. Philos Trans R Soc Lond B Biol Sci, 2009. 364(1526): p. 2063–78.

13. Hines, E.P., et al., Concentrations of phthalate metabolites in milk, urine, saliva, and Serum of lactating North Carolina women. Environ Health Perspect, 2009. 117(1): p. 86–92.

14. Hogberg, J., et al., Phthalate diesters and their metabolites in human breast milk, blood or serum, and urine as biomarkers of exposure in vulnerable populations. Environ Health Perspect, 2008. 116(3): p. 334–9.

15. Du, Y.Y., et al., Follicular fluid and urinary concentrations of phthalate metabolites among infertile women and associations with in vitro fertilization parameters. Reprod Toxicol, 2016. 61: p. 142–50.

16. Pia Dima, A., et al., Development and validation of a liquid chromatography-tandem mass spectrometry method for the simultaneous determination of phthalates and bisphenol a in serum, urine and follicular fluid. Clin Mass Spectrom, 2020. 18: p. 54–65.

17. Johns, L.E., et al., Exposure assessment issues in epidemiology studies of phthalates. Environ Int, 2015. 85: p. 27–39.

18. Hannon, P.R., et al., Mono(2-ethylhexyl) phthalate accelerates early folliculogenesis and inhibits steroidogenesis in cultured mouse whole ovaries and antral follicles. Biol Reprod, 2015. 92(5): p. 120.

19. Wang, W., et al., Mono-(2-ethylhexyl) phthalate induces oxidative stress and inhibits growth of mouse ovarian antral follicles. Biol Reprod, 2012. 87(6): p. 152.

20. Hannon, P.R., et al., Di(2-ethylhexyl) phthalate inhibits antral follicle growth, induces atresia, and inhibits steroid hormone production in cultured mouse antral follicles. Toxicol Appl Pharmacol, 2015. 284(1): p. 42–53.

21. Gore, A.C., et al., EDC-2: The Endocrine Society’s Second Scientific Statement on Endocrine-Disrupting Chemicals. Endocr Rev, 2015. 36(6): p. E1–E150.

22. Land, K.L., et al., The effects of endocrine-disrupting chemicals on ovarian– and ovulation-related fertility outcomes. Mol Reprod Dev, 2022. 89(12): p. 608–631.

23. Hannon, P.R. and J.A. Flaws, The effects of phthalates on the ovary. Front Endocrinol (Lausanne), 2015. 6: p. 8.

24. Cuenca, L., et al., Environmentally-relevant exposure to diethylhexyl phthalate (DEHP) alters regulation of double-strand break formation and crossover designation leading to germline dysfunction in Caenorhabditis elegans. PLoS Genet, 2020. 16(1): p. e1008529.

25. Wang, W., et al., Di (2-ethylhexyl) phthalate inhibits growth of mouse ovarian antral follicles through an oxidative stress pathway. Toxicol Appl Pharmacol, 2012. 258(2): p. 288–95.

26. Liu, J.C., et al., Di (2-ethylhexyl) phthalate impairs primordial follicle assembly by increasing PDE3A expression in oocytes. Environ Pollut, 2021. 270: p. 116088.

27. Tu, Z., et al., Dibutyl phthalate exposure disrupts the progression of meiotic prophase I by interfering with homologous recombination in fetal mouse oocytes. Environ Pollut, 2019. 252(Pt A): p. 388–398.

28. Zhou, C. and J.A. Flaws, Effects of an Environmentally Relevant Phthalate Mixture on Cultured Mouse Antral Follicles. Toxicol Sci, 2017. 156(1): p. 217–229.

29. Land, K.L., et al., Ovulation is Inhibited by an Environmentally Relevant Phthalate Mixture in Mouse Antral Follicles In Vitro. Toxicol Sci, 2021. 179(2): p. 195–205.

30. Fletcher, E.J., et al., Neonatal exposure to an environmentally relevant phthalate mixture alters ovarian function in mice. Toxicol Appl Pharmacol, 2025. 500: p. 117372.

31. Warner, G.R., et al., Environmentally relevant mixtures of phthalates and phthalate metabolites differentially alter the cell cycle and apoptosis in mouse neonatal ovariesdagger. Biol Reprod, 2021. 104(4): p. 806–817.

32. Yao, W., et al., Associations between Phthalate Metabolite Concentrations in Follicular Fluid and Reproductive Outcomes among Women Undergoing in Vitro Fertilization/Intracytoplasmic Sperm Injection Treatment. Environ Health Perspect, 2023. 131(12): p. 127019.

33. Beck, A.L., et al., Ovarian follicular fluid levels of phthalates and benzophenones in relation to fertility outcomes. Environ Int, 2024. 183: p. 108383.

34. Basso, C.G., et al., Associations between urinary and follicular fluid concentrations of phthalate metabolites and reproductive outcomes in Brazilian women undergoing fertility treatment. Reprod Toxicol, 2025. 133: p. 108868.

35. Hoffmann-Dishon, N., et al., Endocrine-disrupting chemical concentrations in follicular fluid and follicular reproductive hormone levels. J Assist Reprod Genet, 2024. 41(6): p. 1637–1642.

36. Yazdy, M.M., et al., A possible approach to improving the reproducibility of urinary concentrations of phthalate metabolites and phenols during pregnancy. J Expo Sci Environ Epidemiol, 2018. 28(5): p. 448–460.

37. Wang, Y., H. Zhu, and K. Kannan, A review of biomonitoring of phthalate exposures. Toxics, 2019. 7(2): p. 21.

38. Warner, G.R., et al., Year-to-year variation in phthalate metabolites in the Midlife Women’s Health Study. Journal of exposure science & environmental epidemiology, 2024. 34(4): p. 610–619.

39. Vogel, N., et al., Current exposure to phthalates and DINCH in European children and adolescents–Results from the HBM4EU Aligned Studies 2014 to 2021. International Journal of Hygiene and Environmental Health, 2023. 249: p. 114101.

40. Koch, H.M. and A.M. Calafat, Human body burdens of chemicals used in plastic manufacture. Philosophical Transactions of the Royal Society B: Biological Sciences, 2009. 364(1526): p. 2063–2078.

41. Gokyer, D., et al., Phthalates are detected in the follicular fluid of adolescents and oocyte donors with associated changes in the cumulus cell transcriptome. F&S Science, 2025. 6(1): p. 30–41.

42. Meling, D.D., et al., The effects of a phthalate metabolite mixture on antral follicle growth and sex steroid synthesis in mice. Toxicol Appl Pharmacol, 2020. 388: p. 114875.

43. Neff, A.M., et al., The role of the aryl hydrocarbon receptor in mediating the effects of mono(2-ethylhexyl) phthalate in mouse ovarian antral folliclesdagger. Biol Reprod, 2024. 110(3): p. 632–641.

44. Yao, Y.C., et al., Predictors of phthalate metabolites in urine and follicular fluid and correlations between urine and follicular fluid phthalate metabolite concentrations among women undergoing in vitro fertilization. Environ Res, 2020. 184: p. 109295.

45. Jiao, X., et al., Iodoacetic acid disrupts mouse oocyte maturation by inducing oxidative stress and spindle abnormalities. Environmental Pollution, 2021. 268: p. 115601.

46. Hagemann-Jensen, M., et al., Single-cell RNA counting at allele and isoform resolution using Smart-seq3. Nat Biotechnol, 2020. 38(6): p. 708–714.

47. Liu, J.C., et al., Di (2-ethylhexyl) phthalate exposure impairs meiotic progression and DNA damage repair in fetal mouse oocytes in vitro. Cell Death Dis, 2017. 8(8): p. e2966.

48. Wang, R., et al., Exposure to diisononyl phthalate deteriorates the quality of porcine oocytes by inducing the apoptosis. Ecotoxicol Environ Saf, 2023. 254: p. 114768.

49. Gu, Y., et al., Exposure to phthalates DEHP and DINP May lead to oxidative damage and lipidomic disruptions in mouse kidney. Chemosphere, 2021. 271: p. 129740.

50. Jabůrek, M., et al., Antioxidant synergy of mitochondrial phospholipase PNPLA8/iPLA2γ with fatty acid–conducting SLC25 gene family transporters. Antioxidants, 2021. 10(5): p. 678.

51. Hannon, P.R. and J.A. Flaws, The effects of phthalates on the ovary. Frontiers in endocrinology, 2015. 6: p. 8.

52. Dalman, A., et al., Effect of mono-(2-ethylhexyl) phthalate (MEHP) on resumption of meiosis, in vitro maturation and embryo development of immature mouse oocytes. Biofactors, 2008. 33(2): p. 149–55.

53. Kalo, D., et al., Carryover Effects of Acute DEHP Exposure on Ovarian Function and Oocyte Developmental Competence in Lactating Cows. PLoS One, 2015. 10(7): p. e0130896.

54. Mu, X., et al., DEHP exposure impairs mouse oocyte cyst breakdown and primordial follicle assembly through estrogen receptor-dependent and independent mechanisms. J Hazard Mater, 2015. 298: p. 232–40.

55. Li, F.P., et al., Di(n-butyl) phthalate exposure impairs meiotic competence and development of mouse oocyte. Environ Pollut, 2019. 246: p. 597–607.

56. Gao, L., et al., Glycine ameliorates MBP-induced meiotic abnormalities and apoptosis by regulating mitochondrial-endoplasmic reticulum interactions in porcine oocytes. Environ Pollut, 2022. 309: p. 119756.

57. Frederiksen, H., et al., Urinary excretion of phthalate metabolites in 129 healthy Danish children and adolescents: estimation of daily phthalate intake. Environ Res, 2011. 111(5): p. 656–63.

58. Henriksen, L.S., et al., Use of stored serum in the study of time trends and geographical differences in exposure of pregnant women to phthalates. Environ Res, 2020. 184: p. 109231.

59. Krotz, S.P., et al., Phthalates and bisphenol do not accumulate in human follicular fluid. J Assist Reprod Genet, 2012. 29(8): p. 773–7.

60. Sun, X., et al., Smart RNA Sequencing Reveals the Toxicological Effects of Diisobutyl Phthalate (DiBP) in Porcine Oocytes. Environ Sci Technol, 2024.

61. Meyer, J.N., et al., Mitochondria as a target of environmental toxicants. Toxicol Sci, 2013. 134(1): p. 1–17.

62. Wang, L.Y., et al., Mitochondrial functions on oocytes and preimplantation embryos. J Zhejiang Univ Sci B, 2009. 10(7): p. 483–92.

