## Supplemental Figures for "Effects of Phthalate Metabolite Mixture Exposure on Mouse Oocyte Development"

**Supplementary Table S1. Purchasing information of each phthalate used in the phthalate mixture.**

| **Abbreviation** | **Full name** | **Company** | **Catalog number** |
| --- | --- | --- | --- |
| MEHP | rac Mono(ethylhexyl) Phthalate | LGC standards | TRC-M542490-100MG |
| MEP | Monoethyl-Phthalate | LGC standards | TRC-M542580-50MG |
| MBP | Mono-Butyl phthalate | Sigma Aldrich | 30751-100MG |
| MiNP | 1,2-Benzenedicarboxylic Acid 1-(7-Methyloctyl) Ester | LGC standards | TRC-B185500-50MG |
| MBzP | Monobenzyl-Phthalate | LGC standards | TRC-M524900-50MG |
| MiBP | Monoisobutyl-Phthalate | LGC standards | TRC-M547700-50MG |

**Supplementary Table S2. The amount of each phthalate used in the phthalate mixture.**

| **Phthalates** | **Percentage** | **Weights (mg/ml)** |
| --- | --- | --- |
| MEHP | 19.4% | 2.878 |
| MEP | 36.7% | 3.798 |
| MBP | 15.3% | 1.811 |
| MiNP | 10.2% | 1.591 |
| MBzP | 8.2% | 1.12 |
| MiBP | 10.2% | 1.209 |

**Supplementary Figure 1**

**Figure S1. High concentrations of phthalate mixture do not disrupt first polar body extrusion but induce chromosome misalignment in MII.**

(A) Representative morphology of oocytes after 18 h of in vitro culture in the control and phthalate-treated groups. Scale bar, 50 μm.

(B) Quantitative analysis of oocytes with first polar body extrusion (PBE) rate in control and phthalate-treated groups after 18 hours of culture. Data in Boxplots show the median, first and third quartiles, and whiskers indicate the range, n=4. ns, not significant (p ≥ 0.05) by two-tailed Student’s t-test.

(C) Immunofluorescence staining of tubulin (green) in control and phthalate-treated groups after 18 hours of culture. Nuclei are counterstained with DAPI (blue). Scale bar: 20 μm.

(D) Proportion of abnormal MII oocytes in the control and phthalate-treated groups after 18 hours of culture. Boxplot representation as in (B), n=4. **p* < 0.05, ***p* < 0.01 by two-tailed Student’s t-test.

**Supplementary Figure 2**

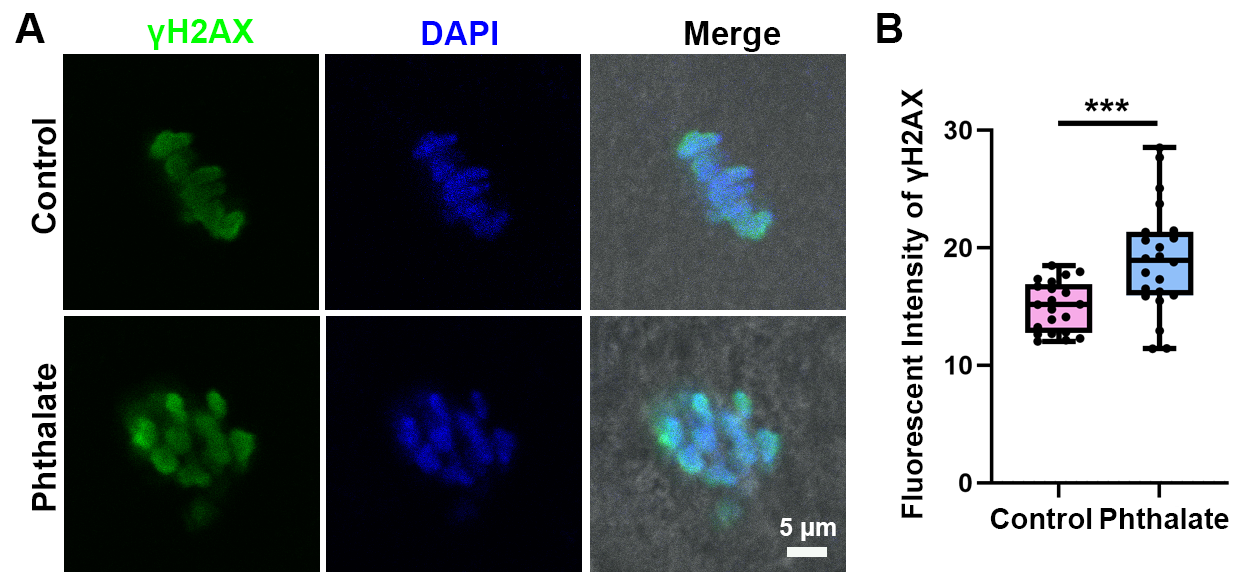
**Figure S2. Phthalate metabolite mixture leads to DNA damage in mouse oocytes.**

(A) Immunofluorescence staining of γ-H2AX (green), a marker of DNA double-strand breaks, in control and phthalate-injected oocytes after 8-hour *in vitro* culture. DNA were counterstained with DAPI (blue). Scale bar: 5 μm.

(B) Quantitative analysis of average γ-H2AX (green) intensity in control and phthalate metabolite mixture-injected oocytes. A total of 21 control oocytes and 22 treated oocytes were analyzed. Data are represented as mean ± SEM of at least three independent experiments, ****P* < 0.001, compared with control.
